# Pulses of melanopsin-directed contrast produce highly reproducible pupil responses that are insensitive to a change in background radiance

**DOI:** 10.1101/365718

**Authors:** Harrison McAdams, Aleksandra Sasha Igdalova, Manuel Spitschan, David H. Brainard, Geoffrey K. Aguirre

## Abstract

**Purpose:** To measure the pupil response to pulses of melanopsin-directed contrast, and compare this response to those evoked by cone-directed contrast and spectrally-narrowband stimuli.

**Methods:** 3-second unipolar pulses were used to elicit pupil responses in human subjects across 3 sessions. Thirty subjects were studied in Session 1, and most returned for Sessions 2 and 3. The stimuli of primary interest were “silent substitution” cone‐ and melanopsin-directed modulations. Red and blue narrowband pulses delivered using the post-illumination pupil response (PIPR) paradigm were also studied. Sessions 1 and 2 were identical, while Session 3 involved modulations around higher radiance backgrounds. The pupil responses were fit by a model whose parameters described response amplitude and temporal shape.

**Results:** Group average pupil responses for all stimuli overlapped extensively across Sessions 1 and 2, indicating high reproducibility. Model fits indicate that the response to melanopsin-directed contrast is prolonged relative to that elicited by cone-directed contrast. The group average cone‐ and melanopsin-directed pupil responses from Session 3 were highly similar to those from Sessions 1 and 2, suggesting that these responses are insensitive to background radiance over the range studied. The increase in radiance enhanced persistent pupil constriction to blue light.

**Conclusions:** The group average pupil response to stimuli designed through silent substitution provides a reliable probe of the function of a melanopsin-mediated system in humans. As disruption of the melanopsin system may relate to clinical pathology, the reproducibility of response suggests that silent substitution pupillometry can test if melanopsin signals differ between clinical groups.

## Introduction

Melanopsin is a photopigment found within the intrinsically photosensitive retinal ganglion cells (ipRGCs; Figure 1a). While they represent a small fraction (~1-3%) of the total retinal ganglion cell population^1–4^, ipRGCs are critical for entrainment of circadian rhythm^5,6^, aversive responses to light^7^, light-induced lacrimation^8^, and control of pupil diameter^9–11^. Disruption of these reflexive visual functions is seen in many clinical conditions, leading to the speculation that dysfunction in the melanopsin system is responsible^7,12–17^. Consequently there is interest in measuring, in humans, a signal that reflects melanopsin function and testing if this signal varies between groups.

**1.**
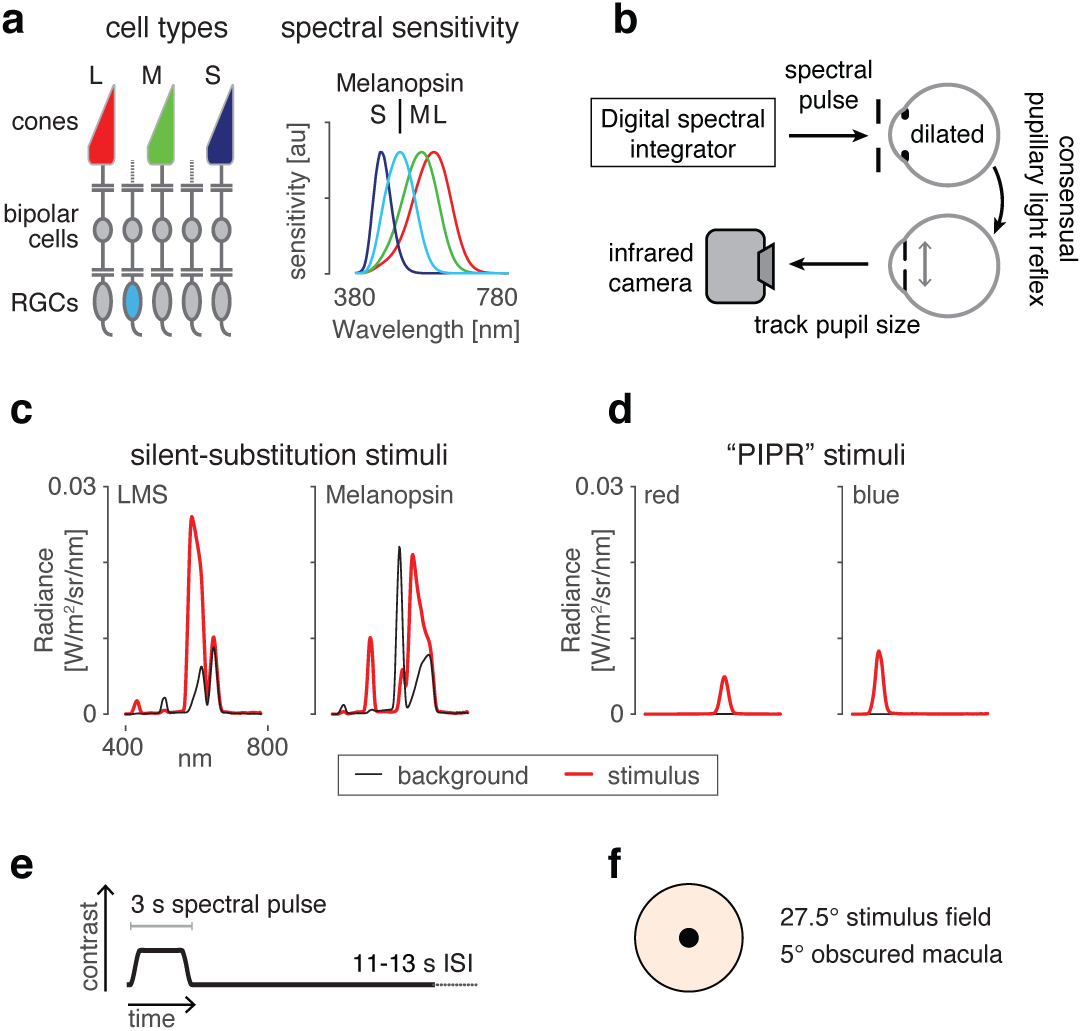
Overview and experimental design. (a, left) L, M, and S cones as well as melanopsin-containing intrinsically sensitive retinal ganglion cells (ipRGCs, blue) mediate visual function at daytime light levels. Although not depicted, ipRGCs receive synaptic input from all three classes of cones. (a, right) The spectral sensitivity functions of these photoreceptors. (b) A digital light integrator delivers spectral pulses to the pharmacologically dilated right eye of the subject’s pupil. The consensual pupillary reflex from the left eye is recorded via an infrared camera. (c) We use silent substitution to selectively target either the L, M and S cones and thus the postreceptoral luminance channel left) or melanopsin right). d) The PIPR stimuli consist of narrowband pulses of long wavelength red light (left) or short wavelength blue light (right). Note that the stimuli are equated in terms of retinal irradiance expressed in quantal units, but because the number of quanta/Watt and pre-receptoral filtering are wavelength dependent, the blue stimulus has higher radiance. All stimuli are from Session 1. The particular spectra plotted here and in panel d are an example from one subject; the spectra varied by the age of the subject to account for preceptoral filtering. (e) We delivered 3 s spectral pulses smoothed by a 500 ms half-cosine window, with an inter-stimulus interval between trials ranging from 11 to 13 s. (f) Stimuli were presented through an eyepiece with a 27.5° with the central 5° obscured to prevent activation of the macula. Some panels are adapted from a prior publication^42^.

The post-illumination pupil response (PIPR) paradigm is one method to assess melanopsin function in humans^11,18,19^. The PIPR paradigm exploits the differing spectral sensitivities of the melanopsin photopigment and the cone-based luminance mechanism: the medium and long-wavelength cones (M and L)—which are the primary input to the luminance mechanism—are more sensitive to light of longer wavelengths (‘red’), while melanopsin sensitivity is greatest in the short-wavelength (‘blue’) range. The PIPR paradigm measures the response of the pupil to pulses of narrowband blue and red light presented against steady dark backgrounds. Particular attention is paid to the behavior of the pupil at relatively delayed time periods, including after stimulus offset i.e., ‘post-illumination’), when melanopsin is found to exert greater influence over pupil size relative to the cones^11,20^. PIPR measurements have been made in numerous clinical conditions, including multiple sclerosis, Parkinson’s disease, idiopathic intracranial hypertension, traumatic brain injury, glaucoma, diabetes, retinitis pigmentosa, Leber’s hereditary optic neuropathy, Smith-Magenis syndrome, and depression^21–32^.

While relatively simple to deploy and measure, interpretation of the PIPR as a melanopsin-specific signal is less straightforward. Because the blue stimulus is presented against a dark background, the pupil response will include a rod contribution^33,34^. Blue light also drives S-cones, which, like melanopsin, produce delayed and sustained pupil responses^35^. While there is convincing evidence that sustained pupil constriction can be produced by melanopsin alone^11^, cones may also contribute (perhaps via the ipRGCs) to a sustained response^36–38^. Therefore, while the PIPR response reflects perhaps overwhelmingly) the contribution of melanopsin signals, it cannot be concluded that differences between clinical populations in PIPR responses are attributable solely to the melanopsin system.

Silent substitution spectral modulations^39^ provide an alternate approach to the study of the melanopsin contribution to the human pupil response^35,40–42^. Light spectra are tailored to modulate the response of one or more targeted photoreceptor mechanisms (e.g., melanopsin), while holding the response of the remaining photoreceptor mechanisms e.g., L-, M‐ and S-cones) constant. Subjects first adapt to a background light spectrum. When the silent substitution modulation is presented around that background, the subsequent response is attributable to the targeted photoreceptor (s). Here, we measured the temporal properties and reliability of the across-subject average pupil response to pulses of melanopsin contrast delivered via silent substitution. We compared the response to melanopsin stimulation to that evoked by cone-directed contrast that was silent for melanopsin, and by narrowband PIPR stimuli. To anticipate, we find that the silent-substitution approach produces a highly reproducible measure of melanopsin-driven pupil response that is insensitive to a change in background radiance.

These experiments were the subject of pre-registration documents. The preregistered protocol was followed with small exceptions, see Methods) in subject recruitment, screening, exclusion, stimulus validation, and pupil data preprocessing. The analyses described in the pre-registration examined the reliability of between subject differences in response. We found relatively low reliability and present those results in the supplementary materials (Supplementary Figures 3, 4). We focus here upon population level analyses that were not pre-registered.

## Methods

### Subjects

Subjects were recruited from the community of students and staff at the University of Pennsylvania. Exclusion criteria for enrollment included a prior history of glaucoma or a negative reaction to pupil-dilating eye drops. During an initial screening session, subjects were also excluded for abnormal color vision as determined by the Ishihara plates^43^ or visual acuity below 20/40 in each eye as determined using a distance Snellen eye chart. Subjects completed a brief, screening pupillometry session. We excluded at this preliminary stage subjects who were unable to provide high-quality pupil tracking data (details below). Poor data quality was found to result from difficulty suppressing blinks or from poor infra-red contrast between the pupil and iris.

A total of 32 subjects were recruited and completed initial screening. Two of these subjects were excluded after screening due to poor data quality e.g., excessive loss of data points from blinking) as determined by pre-registered criteria. Thirty subjects thus successfully completed Session 1 and provided data for analysis. These subjects were between 19-33 years of age (mean 25.93 ± 4.24 SD). Fourteen subjects identified as male, 15 female, and one declined to provide a gender identification. Of this group of 30 subjects, 24 completed an identical second session of testing and 21 completed a third session at higher light levels. The time between participation in Session 1 and Session 2 was on average 110 days, and between Session 1 and Session3 on average 296 days. The study was approved by the Institutional Review Board of the University of Pennsylvania, with all subjects providing informed written consent, and all experiments adhered to the tenets of the Declaration of Helsinki.

When a subject arrived for a session of primary data collection, the right eye was first anesthetized with 0.5% proparacaine and dilated with 1% tropicamide ophthalmic solution. Subjects then had their right eye dark adapted by wearing swimming goggles with the right eye obscured while sitting in a dark room for 20 minutes. In an attempt to minimize variation in circadian cycle across sessions, testing for Sessions 2 and 3 started within three hours of the time of day when the same subject started Session 1.

### Stimuli

The experiments used two classes of stimuli: 1) silent-substitution spectral modulations that targeted either the melanopsin photopigment or the cone-mediated luminance post-receptoral mechanism; 2) narrow-band blue and red stimuli designed to elicit the post-illumination pupil response (PIPR).

The silent-substitution stimuli were a subset of those used in a prior report^42^, and full details of their generation may be found there. Briefly, we used the method of silent substitution together with a digital light synthesis engine (OneLight Spectra) to stimulate targeted photoreceptors. The device produces stimulus spectra as mixtures of 56 independent primaries (~16 nm FWHM) under digital control, and can modulate between these spectra at 256 Hz. Details regarding the device, stimulus generation, and estimates of precision have been previously reported^35,44,45^. Our estimates of photoreceptor spectral sensitivities were as previously described^45^, with those for the cones following the CIE physiological cone fundamentals^46^. The estimates account for the size of the visual field (27.5° diameter), subject age, and the pupil size (which was fixed at 6 mm diameter through the use of an artificial pupil). Separate background and modulation spectra were identified to provide nominal 400% Weber contrast on melanopsin while silencing the cones for the melanopsin-directed background/modulation pair (Mel), and 400% contrast on each of the L-, M-, and S-cone classes while silencing melanopsin for the luminance-directed modulation/background pair (LMS) (Figure 1c). The calculated CIE 1931 chromaticities of the background spectra for the Mel and LMS stimuli were similar (Mel: ~0.55, ~0.41; LMS: ~0.57, ~0.38)^47^. The background for Mel and LMS pulses were nominally rod-saturating (~100 cd/m^2^ for Mel and ~40 cd/m^2^ for LMS for Sessions 1 and 2; ~270 cd/m^2^ and ~100 cd/m^2^ for Session 3). The modulations did not explicitly silence rods or penumbral cones^45^.

The PIPR stimuli consisted of narrowband pulses of blue (475 ± 25 nm peak ± Gaussian FWHM) and red (623 ± 25 nm) light (Figure 1d). These stimuli were each designed to produce 12.30 log quanta-cm^−2^-sec^−1^ retinal irradiance for Sessions 1 and 2, and12.85 log quanta-cm^−2^-sec^−1^ for Session 3, in a manner that accounted for differences in lens density due to subject age. These stimuli were presented against a dim background (~0.5 cd/m^2^ for the first 2 sessions, ~1 cd/m^2^ for the third session). The irradiance of the PIPR stimuli was limited by the gamut of the device at short wavelengths, and the requirement to match the retinal irradiance of the red and blue stimuli. Background light levels were the minimum possible with our apparatus, as some light is emitted by the light engine even when all primaries are set to their minimum level.

Due to imperfections in device control, the actual stimuli presented differed in photoreceptor contrast and irradiance from their nominal designed values.

Before and after each subject’s measurement session, spectroradiometric validation measurements of the background and modulation spectra were obtained. From these, we calculated the actual contrast upon targeted and nominally silenced photoreceptors for that subject (using age-based photoreceptor sensitivities) for the silent substitution stimuli, as well as the retinal irradiance of the PIPR stimuli. Following our pre-registered protocol, we excluded data for a given session if the post-experiment validation measurements showed that the silent substitution stimuli were of insufficient quality. Specifically, if the contrast on the targeted post-receptoral mechanism (Mel or LMS) was less than 350% (as compared to the nominal 400%), or if contrast upon an ostensibly silenced post-receptoral mechanism (Mel, LMS, L–M, or S) was greater than 20%. Data from five sessions were discarded (and subsequently recollected) as a consequence of this procedure. We did not evaluate the PIPR stimuli for the purposes of data exclusion. Supplementary Table 1 provides the results of the stimulus validations for all subjects, sessions, and stimuli. These calculations do not account for the biological variability in individual photoreceptor spectral sensitivity that can produce further departures from nominal stimulus contrasts^42^.

Three-second pulses of spectral change were presented during individual trials of 17 s duration (Figure 1e). During each trial, a transition from the background to the stimulation spectrum (Mel, LMS, blue, or red) would occur starting at either 0, 1, or 2 seconds after trial onset (randomized uniformly across trials); this jitter was designed to reduce the ability of the subject to anticipate the moment of stimulus onset. The transition from the background to the stimulation spectrum, and the return to background, was smoothed by a 500 msec half-cosine window. The half-cosine windowing of the stimulus was designed to minimize perception of a Purkinje tree percept in the melanopsin directed stimulus^45^.

Each session consisted of three blocks of stimuli: PIPR (consisting of both red and blue stimuli counterbalanced in order within subject), LMS, and Mel, in this fixed order. At the start of each block the subject adapted to the background spectrum for 4.5 minutes. The block consisted of twenty-four, 17 second trials. Within each block, after every 6 trials, participants were permitted to take a break before resuming the experiment.

Stimuli were presented through a custom-made eyepiece with a circular, uniform field of 27.5° diameter and the central 5° diameter obscured (Figure 1f).

The central area of the stimulus was obscured to minimize stimulation within the macula, where macular pigment alters the spectral properties of the stimulus arriving at the photoreceptors. Subjects viewed the field through a 6 mm diameter artificial pupil and were asked to maintain fixation on the center of the obscured central region.

### Pupillometry

Pupil diameter was measured using an infrared video pupillometry system (Video Eye Tracker; Cambridge Research Systems Ltd.), sampled at 50 Hz. Following acquisition, the raw measured pupil response was adjusted in time to account for the stimulus onset time within each trial and normalized by the baseline pupil size for that trial (with baseline size taken as the mean pupil diameter for 1 second prior to stimulus onset). Data points in the resulting response for which the velocity of constriction or dilation exceeded 2500% change/s) were rejected and replaced via linear interpolation. The responses across trials were averaged.

Pupillometry data were excluded from analysis on the basis of the number of rejected data points. Trials containing 10% or more rejected data points were deemed incomplete and excluded from the average; if more than 75% of the trials for a given stimulus type were excluded, then the entire session was judged to be incomplete and the subject was either re-studied or excluded, following our preregistered procedure. Additionally, if more than 50% of trials across all stimulus types were excluded, then the subject was either re-studied or excluded. Data from four sessions were discarded for this reason; 2 of these subjects were re-studied.

As noted briefly under Subjects above, screening pupillometry was also performed prior to primary data collection to exclude subjects for whom good quality pupil tracking data could not be obtained. In a screening session, subjects were presented two sets of six trials of the PIPR stimuli. Subjects with 4 or more incomplete trials assessed by the same criterion above were excused from the experiment. Two subjects were excused from the study in this manner.

### Analysis

We fit the pupil response for each stimulus and subject using a three-component temporal model (Figure 3a)^42^. The stimulus profile passes through the model and, under the control of six parameters, is transformed into a predicted pupil response. The six parameters include two time constants that influence the shape of each component, three gain parameters that adjust the scaling of each component, and one onset delay parameter that shifts the entire modeled response in time. The transient component captures the initial peak of pupil constriction, the sustained component tracks the shape of the stimulus profile, and the persistent component describes the slow dilation of the pupil back to baseline. Each component has an amplitude parameter. The shape of the components are under the control of two temporal parameters. The τ_gamma_ parameter controls the rate of onset and width of all components. The τ_exponential_ controls the rate of exponential decay of the persistent component. The three components are summed to create the model response, which is then temporally shifted in time by the overall delay parameter. We fit this model to the average response for each subject for each stimulus condition. In analyzing group differences of model parameters, the median value was used as parameters were not normally distributed across subjects.

**3.**
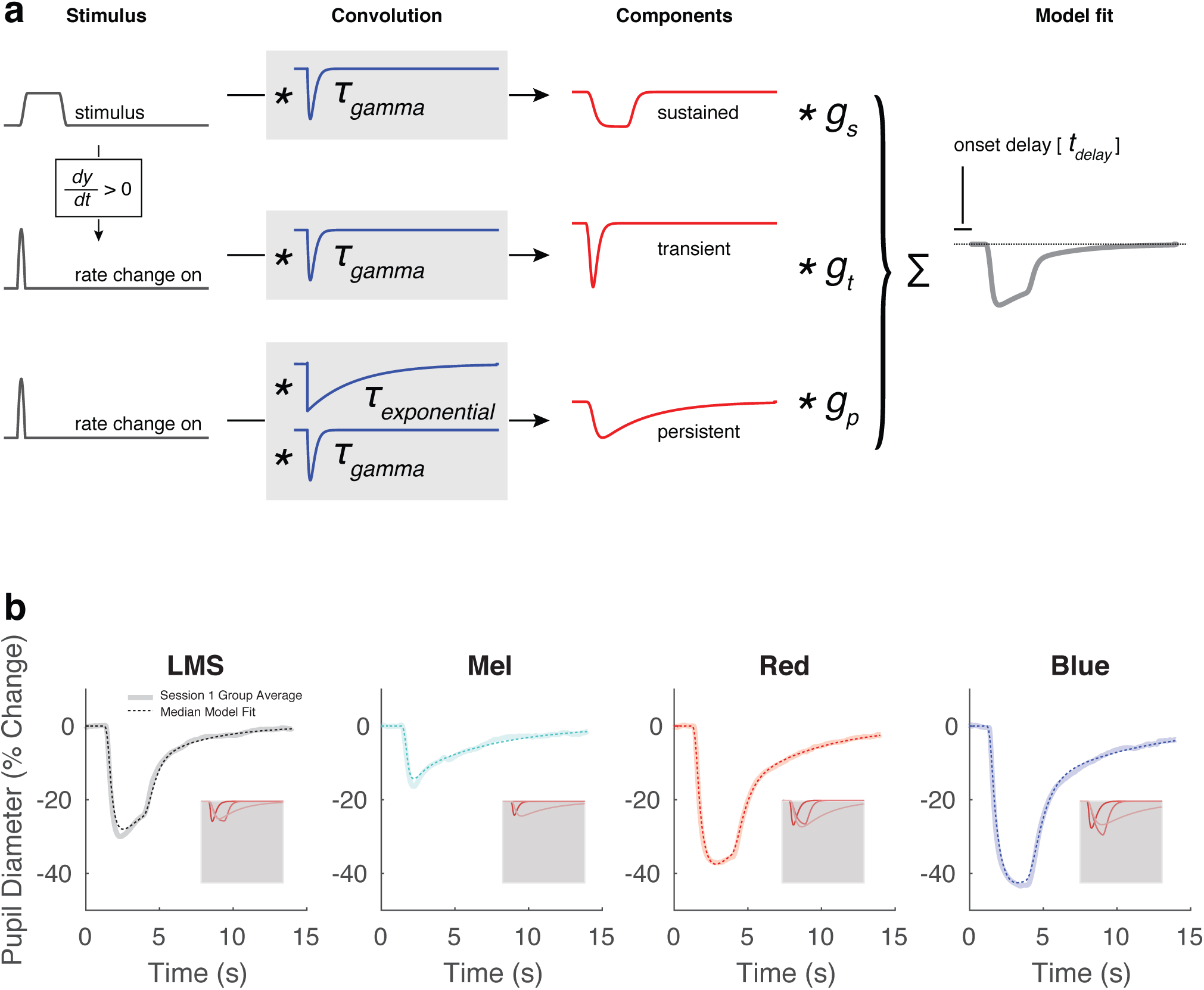
A three-component model used to fit the group average pupil responses. (a) Within-subject average evoked responses to each stimulus type was subjected to non-linear fitting with a six-parameter, three-component model. The model was designed to capture the visually apparent and temporally separated components of the evoked pupil response. The elements of the model are not intended to directly correspond to any particular biological mechanism. The input to the model was the stimulus profile (black). An additional input vector, representing the rate of stimulus change at onset, was created by differentiating the stimulus profile and retaining the positive elements. These three vectors were then subjected to convolution operations composed of a gamma and exponential decay function (blue), each under the control of a single time-constant parameter (τ _gamma_ and τ _exponential_). The resulting three components (red) were normalized to have unit area, and then subjected to multiplicative scaling by a gain parameter applied to each component (g_transient_, g_sustained_, and g_persistent_). The scaled components were summed to produce the modeled response (gray), which was temporally shifted (t_delay_). This caption and the corresponding panel are adapted from Figure S9 of Spitschan et al. 2017^42^. (b) The model fit, computed from the median response parameter across all 30 subjects from Session 1, is plotted in dotted lines on top of the group average response from Session 1. The gray inset shows each model component of the fit (transient, sustained, and persistent in most to least saturated color).

Model fits were performed using MATLAB’s fmincon function. Fits were initialized from 6 different starting positions and the fit with the highest proportion variance explained (R^2^) was retained. Additionally, bounds were placed on each parameter, as informed by an initial inspection of the data. The bounds of the τ_gamma_ were different for responses elicited through silent substitution and PIPR stimuli. Specifically, the upper boundary of τ_gamma_ for fits to responses elicited by PIPR stimuli was greater than that for fits to responses elicited through silent substitution to reflect the generally wider shape of these responses. This choice improved the quality of fits to each stimulus type. As we were interested exclusively in comparisons within a stimulus type (LMS vs. Mel, red vs. blue), the differing parameter boundaries would not influence any subsequent conclusions. We also performed additional analyses in which we locked and freed different sets of parameters as part of control tests. These procedures are described in the Supplementary Materials (Supplementary Figure 1).

To test for significance of observed group differences of metrics derived from our model, we used label permutation. For a given group comparison, we took the observed metric aggregated across all trials for a given stimulus type for each subject and randomly assigned each metric to the correct stimulus label or the opposite stimulus label. After performing this for all subjects, we computed the median difference. We performed this simulation 1,000,000 times, and asked the percentage of simulations in which the simulated median difference is more extreme than the observed median difference.

### Pre-registration of studies

Our studies (composed of three sessions of data collection) were the subject of preregistration documents (https://osf.io/9umq4/) and annotated addenda (https://osf.io/bg76w/). The pre-registered protocol dictated subject recruitment, screening, data exclusion, and stimulus validation. Session 1 was designed to test if we could measure the melanopsin mediated pupil response to silent substitution and PIPR stimuli in individuals. Data collection for Session 1 commenced in September 2016. An addendum (https://osf.io/hyj89/) detailed an improvement in our approach to generating stimuli that accurately described stimulus production for both the initial and subsequent subjects; this document is dated September 2016 but was not uploaded until October 2016. A January of 2017 addendum clarified an ambiguity in our original description of the stimulus validation procedure (https://osf.io/b4r3q/).

Session 2 was designed to determine if the magnitude of pupil response to melanopsin stimulation was a reliable individual subject difference (https://osf.io/z2vj7/). Session 3 repeated the measurements at a higher light level in an attempt to evoke a larger response to the PIPR stimuli and to further test the reliability of any individual differences in pupil response (https://osf.io/angyu/).

Our original motivation for these studies was to measure individual differences in pupil response. We ultimately determined that this test was limited by within-session measurement noise. Therefore, this paper focuses on comparisons at the group level. In keeping with our pre-registered protocols, however, we provide in the supplementary material the results of individual subject analyses (Supplementary Figures 3, 4).

There was ambiguity in our initial protocol regarding the interpretation of post-experimental stimulus validations. Five validation measurements were made after each experimental session. Our pre-registration initially failed to delineate how to interpret all five validation values; our procedure was later clarified to specify that data from a session would be excluded if the median value across all five post-experiment validation values was larger than the cutoff criterion. Data from one session were discarded and re-collected based upon an initial interpretation of the validation procedure in which a single validation measurement that exceeded criterion led to data rejection. Following the clarification of our procedure to use the median validation measurement, data from four subsequent sessions were discarded and recollected because of stimulus quality.

### Availability of data and analysis code

Data will be available via figshare upon publication. Analysis code that operates upon the raw data and produces the results and figures may be found here: https://github.com/gkaguirrelab/pupilPIPRAnalysis.

## Results

In each of 30 subjects we measured consensual pupil responses in the left eye to spectral modulations presented to the pharmacologically dilated right eye (Fig 1b). Two of the modulations targeted the post-receptoral luminance or melanopsin pathway using a silent-substitution spectral exchange (Fig 1c, left), and two of the modulations were narrowband red or blue increments typical of PIPR studies (Fig 1c, right). We presented these spectral modulations as half-cosine windowed, 3 second pulses (Fig 1d) on a spatially uniform field, except for masking of the central 5° of visual angle to minimize stimulation of the macula (Fig 1e). We recorded the ensuing pupil response for each of many trials in 30 subjects (Session 1) and a subset of these subjects in Sessions 2 (24 subjects) and 3 (21 subjects).

### Silent substitution and PIPR stimuli elicit highly reproducible pupil responses at the group level

We first examined the form of group (averaged over subjects) pupil responses to pulsed spectral modulations designed to selectively target the cones or melanopsin (Figure 2a, top row). We measured pupil responses during the 13 seconds that followed the onset of a 3 second stimulus pulse, and expressed pupil size as the percentage change in diameter relative to the pre-stimulus period. For our silent-substitution stimuli, which were equated in contrast, the LMS-mediated pupil response was of overall larger amplitude than that evoked by the melanopsin-directed stimulus. The responses also differed in their shape, with the offset of the stimulus producing a more rapid dilation for LMS stimulation as compared to melanopsin stimulation.

**2.**
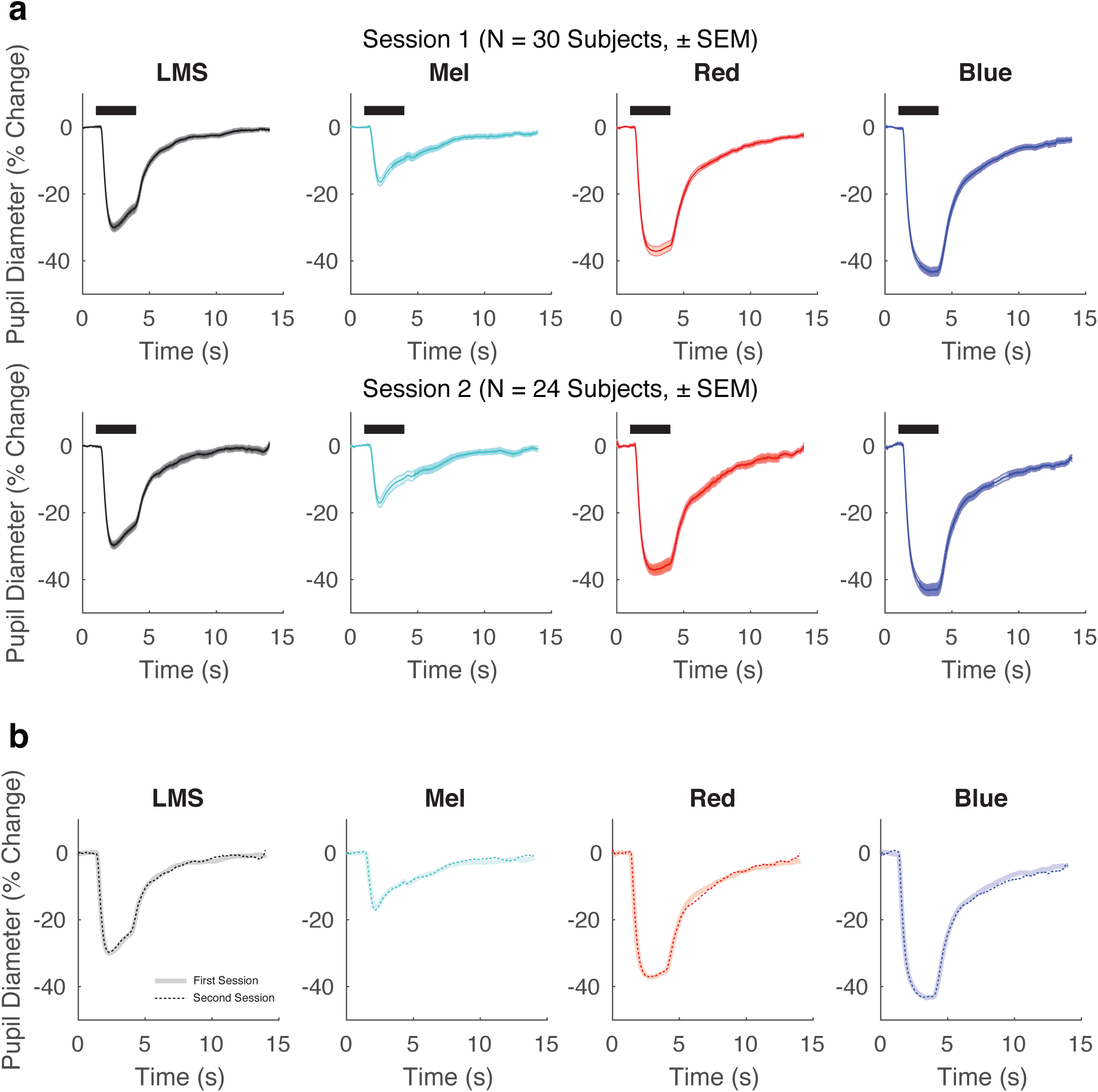
The group average pupil response is stable over time. (a) Group average pupil responses (± standard error of the mean) for each stimulus condition from Session 1 (top, N = 30 subjects) and Session 2 (bottom, N = 24 subjects). (b) Group average responses from Session 2 in thicker, desaturated colors. Responses from Session 1 indicated with thinner, dotted lines. The two lines overlap extensively.

The red and blue PIPR stimuli also produced pupil constriction. These stimuli were equated in retinal irradiance but the amplitude of pupil constriction was smaller in response to the red stimulus as compared to the blue stimulus. The shape of these responses also differed subtly, as the pupil began to dilate during the red stimulus, while the constriction in response to the blue stimulus continued to increase during stimulus presentation.

The standard error of the mean across subjects was quite small relative to the amplitude of response. While this might suggest that the measurements would be reproducible in this group, it is possible that variation in subject state (e.g., due seasonal or circadian changes) or drift in our apparatus would reduce reproducibility across sessions. We tested for reproducibility by repeating the measurements during Session 2 in 24 of the 30 subjects between 54 and 175 days later (Fig 2a, bottom row). The amplitude, shape, and within-session standard error of the mean measured in Session 2 was quite similar to that measured in Session 1. Figure 2b presents the group average from Session 2 plotted directly on top of that from Session 1 for each stimulus condition. The reproducibility of the pupil response to all stimuli is evident, both in amplitude max absolute difference in amplitude of group-averaged responses: 1.47% for LMS, 1.64% for Mel, 2.24% for red 1.74%, for blue), and in shape (Pearson correlation coefficient of the Session 1 group average with the Session 2 group average: r = 0.999 for LMS, r = 0.995 for Mel, 0.998 for red, r = 0.999 for blue).

### The melanopsin response is more persistent than the cone response

Melanopsin driven activation of ipRGCs results in notably prolonged responses^5,20^. Here we asked if a difference in the temporal profile of the pupil response to cone and melanopsin stimulation is apparent at the group level. We fit the data from each subject with a three-component model of the pupil response (Figure 3a)^42^. The model has amplitude parameters for transient, sustained, and persistent components, as well as three temporal parameters that specify the overall timing and influence the shape of the components. Figure 3b illustrates the model fits for the data from Session 1. The fit line is given by the median of the model parameters across subjects. There is good agreement between the model and the across-subject average response. The amplitude and shape of the three model components are shown inset in each panel. After combining the data from Sessions 1 and 2 for those subjects studied twice, we tested for differences in the amplitude and temporal parameters evoked by the different stimuli.

The transient, sustained, and persistent components of the model reflect different temporal domains. The persistent component captures the slow return to baseline of the pupil response following the offset of the stimulus. We considered that stimulation of the ipRGCs might produce pupil responses with a relatively enhanced persistent component. For each subject for each stimulus type, we computed the proportion of the total pupil response area made up of the persistent component (Figure 4a). Across subjects, the median pupil response to LMS stimulation had 50% of its total response area fit by the persistent component. In contrast, the response to melanopsin stimulation was 76% persistent (p = 0.0015 established by permutation of stimulus labels). This difference reflects primarily a larger sustained component in the pupil response to luminance; the absolute response area of the persistent component was similar for the cone and melanopsin driven responses (Supplementary Table 2). Unexpectedly, for the PIPR stimuli, the persistent component was larger in response to the red as compared to the blue stimulus (median ‘percent persistent’ for red: 65%; for blue: 58%; p = 0.00047 by label permutation).

**4.**
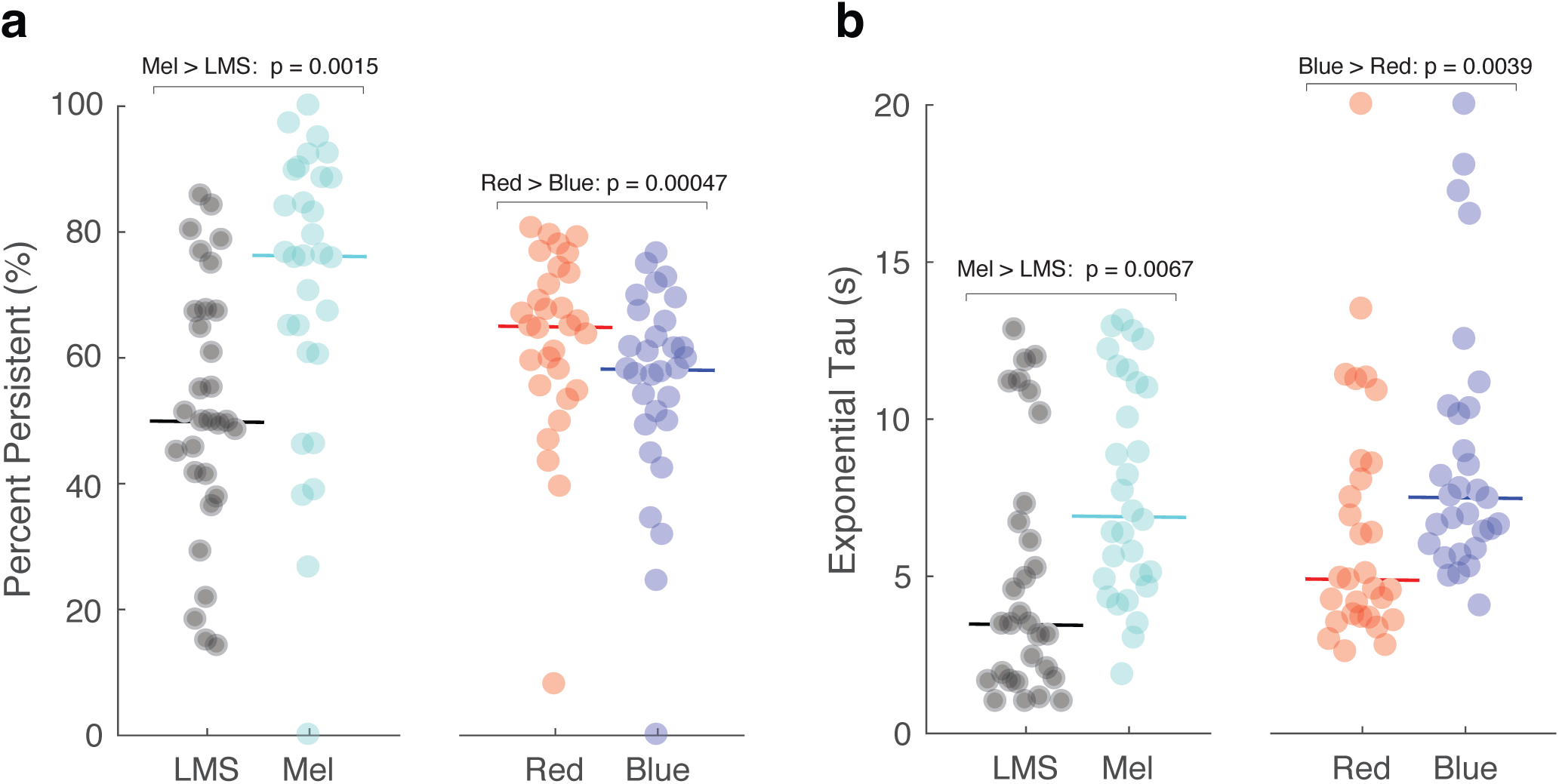
Model parameters derived from the pupil responses to the different stimuli. The pupil response (averaged across trials and across Sessions 1 and 2) was obtained for each subject and stimulus and then fit with the temporal model. (a) The area of the persistent component of the pupil response was scaled by the total response area (N = 30 subjects). The solid horizontal line indicates the median value across subjects. Permutation testing was used to assess the significance of median differences in response across stimulus conditions at the group level. (b) The exponential tau parameter across subjects and stimuli.

We considered that the temporal profile of the persistent response, as opposed to its magnitude alone, would reflect the influence of melanopsin. The model parameter τ_exponential_ influences the rate at which the persistent component of pupil diameter dilates back to baseline following stimulus offset. We tested if this time constant differed in the responses to the stimulus types. Consistent with the expected properties of the ipRGCs, the melanopsin driven response had a slower return to baseline as compared to the LMS-driven response Figure 4b, median τ_exponential_ for LMS: 3.46 s; for Mel: 6.90 s; p = 0.0067 by label permutation). This slower return to baseline was also observed for the response to the blue stimulus as compared to the red stimulus (red: 4.89 s; blue: 7.50 s; p = 0.0039 by label permutation). Supplementary Table 2 contains the amplitude and temporal parameters for all conditions and stimuli.

A property of our analysis is that the temporal parameters are allowed to vary between the compared stimulus conditions to best fit the data. It is therefore possible that observed differences in the τ_exponential_ or ‘percent persistent’ measurements arise as a consequence of differences in other model parameters. To evaluate this possibility, we re-ran the analyses holding the other temporal parameters fixed between the two compared stimulus conditions. This analysis revealed very similar results (Supplementary Figure 1).

### Silent substitution and PIPR methods are differently sensitive to stimulus radiance

We considered the possibility that the pupil response evoked by the silent substitution stimuli would be relatively insensitive to the overall spectral power of the stimuli, as long as the contrast was held constant. For Session 3, we modified our apparatus to increase the radiance of all stimuli. Although the background luminance of the silent substitution stimuli more than doubled (mean background luminance for LMS from ~40 cd/m^2^ to ~100 cd/m^2^; for Mel increased from ~100 cd/m^2^ to ~270 cd/m^2^), the calculated LMS and melanopsin contrast remained the same. For the PIPR stimuli, the nominal intensity of the spectral pulse increased from 12.30 to 12.85 log quanta-cm^−2^-sec^−1^, and the background luminance increased from 0.5 cd/m^2^ to 0.9 cd/m^2^).

We then repeated the pupil measurements in 21 of the 30 subjects between 238 and 352 days after their initial enrollment (Session 3). Figure 5 presents the group average response collapsed across the first two sessions, compared to the pupil response measured in Session 3. For the LMS and the Mel stimuli the average group response was essentially unchanged (max absolute difference in amplitude of group-averaged responses: 1.10% for LMS, 1.27% for Mel; Pearson correlation of the evoked response between Session 1/2 and Session 3: LMS, r = 0.999; Mel, r = 0.994). This high degree of reproducibility suggests that the pupil response to the silent substitution stimuli is insensitive to this change in absolute light intensity and instead reflects stimulus contrast.

**5.**
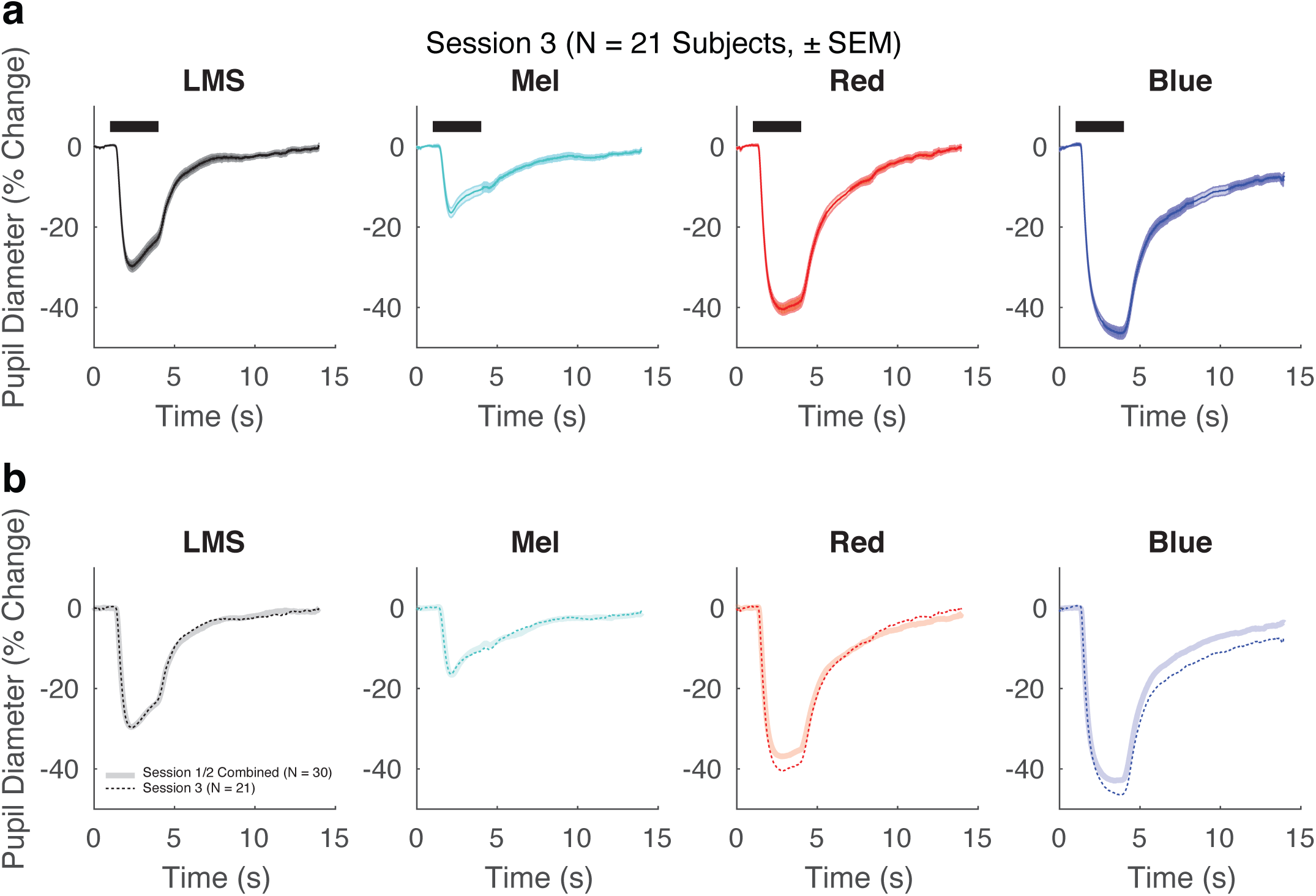
The effect of an increase in stimulus radiance. (a) Group average pupil responses (± standard error of the mean) for each stimulus condition from Session 3 (N = 21 subjects). b) Group average responses from Session 3 background luminance for Mel and LMS were ~270 cd/m2 and ~ 100 cd/m2, respectively) in dotted lines are plotted on top of group average responses from Sessions 1 and 2 combined (N = 30 subjects; background luminance for Mel and LMS were ~100 cd/m2 and ~40 cd/m2, respectively). While the change in stimulus radiance did not alter the pupil response to the silent substitution stimuli, the pupil response to the PIPR stimuli was increased.

In distinction, the increase in the radiance of the PIPR stimuli produced a larger amplitude of pupil response (max absolute difference in amplitude of group-averaged responses: 3.60% for red, 4.89% for blue). Many studies that use the PIPR stimuli attempt to isolate the melanopsin-specific component by taking the difference of the blue and red responses. Figure 6 presents the difference in pupil response evoked by the red and blue stimuli at the two radiance levels. This PIPR effect, especially at the later time points, grows in magnitude from Sessions 1 and 2 to Session 3 (Fig 6). We quantified the PIPR effect as the difference in the total response area of the model fits to blue and red stimuli for each subject, for each session. The median PIPR was larger in Session 3 as stimulus radiance was increased (Sessions 1/2 median PIPR: 35 % change * s; Session 3 median PIPR: 74 % change * s; p = 0.0002 by label permutation).

**6.**
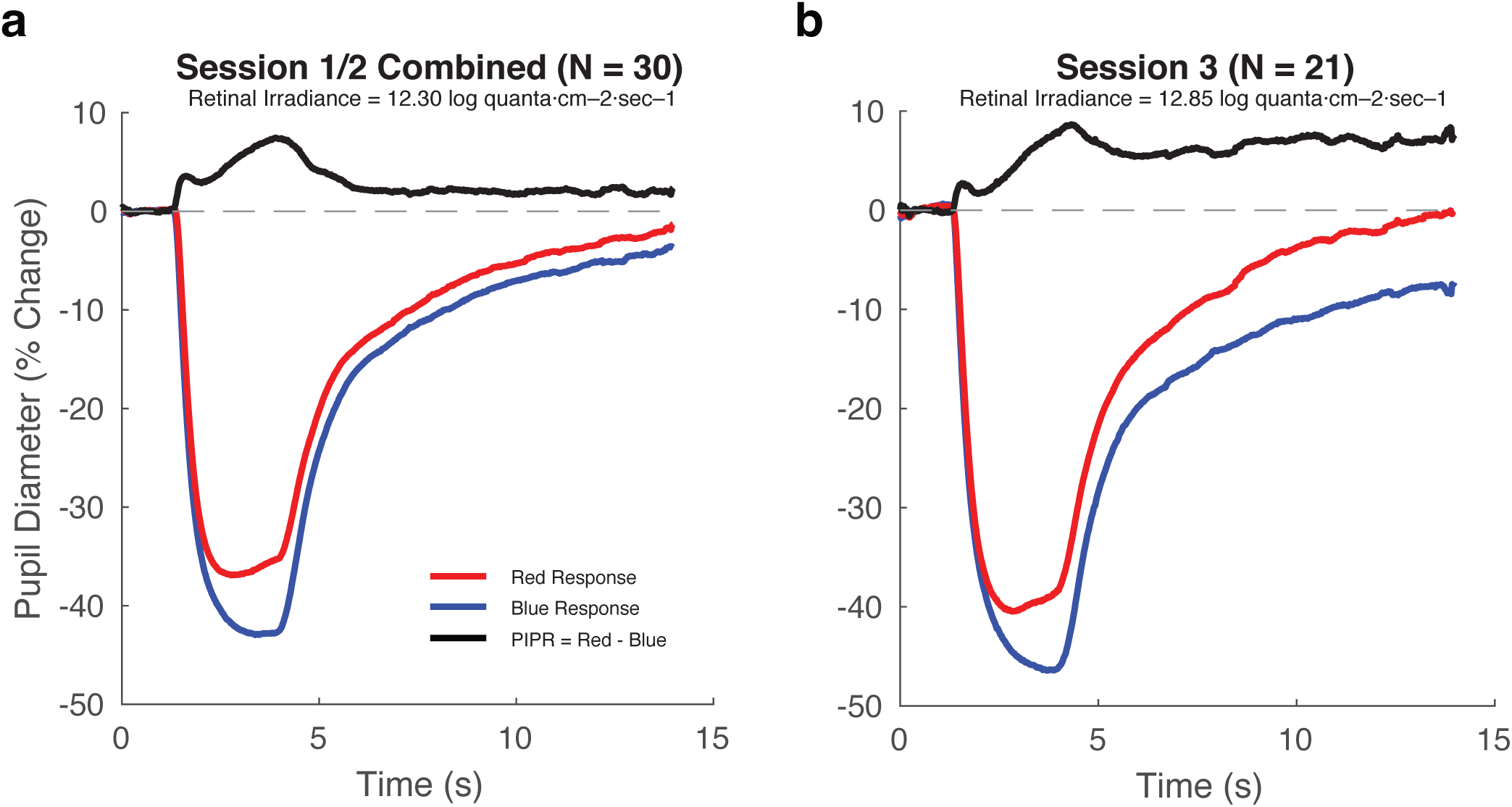
The PIPR effect increases with stimulus intensity. The PIPR effect (black) was obtained by subtracting the blue response from the red response (this order was chosen to provide a positive differential). a) Sessions 1 and 2 presented stimuli with a retinal irradiance of 12.30 log quanta·cm^−2^·sec^−1^ (b). Session 3 used pulses with retinal irradiances of 12.85 log quanta·cm^−2^·sec^−1^.

## Discussion

We find that pulsed spectral modulations that target the cones and melanopsin evoke distinctive pupil responses. At a group level, the average responses to these silent substitution stimuli are highly reliable. Consistent with the known temporal properties of the ipRGCs, the response to melanopsin-directed as compared to cone-directed stimulation features a relatively larger persistent response that returns to baseline more slowly.

Our findings indicate the feasibility of using pupillometry with silent substitution stimuli to test for group differences in cone and melanopsin physiology. As compared to the PIPR stimuli, the silent substitution approach more directly targets and isolates the melanopsin and cone systems. Further, the highly reproducible responses seen at the group level indicate that differences between groups should be detected with good statistical power. Indeed, the extent to which this group average signal is reliable can be seen in the highly similar responses elicited from a different cohort of subjects using the same stimuli as part of a previous study^42^ (Supplementary Figure 2).

We applied a model to the temporal profile of pupil responses to derive amplitude and timing parameters. This model accounts well for the form of response to both silent substitution and PIPR stimuli. Although we use ‘percent persistent’ to describe differences between the cone‐ and melanopsin-driven pupil responses, we find that all stimuli evoke some degree of persistent response. This observation is consistent with prior work, both in previous PIPR studies that show that the red stimulus evokes persistent pupil constriction, as well as neurophysiologic studies that show ipRGCs generate persistent firing from non-melanopsin inputs^38^. We examined as well the τ_exponential_ timing parameter of our model fits. The melanopsin-directed and PIPR blue stimuli produced responses with greater τ_exponential_ values as compared to their cone-directed and PIPR red counterparts. Therefore, while all stimulus types evoked some amount of persistent pupil response, slower resolution of this response was seen for the stimuli thought to drive melanopsin. We anticipate that the temporal model may be used to test for differences in the amplitude and temporal properties of melanopsin-driven responses in clinical populations.

An original motivation for our study was to examine individual differences in the pupil response. While average responses at the group level were highly reliable, we found that there was relatively poor reproducibility for individual subjects (Supplementary Figure 3). We examined the reproducibility of total pupil response amplitude across subjects. While there was a reasonable correlation of this measure between Sessions 1 and 2, these responses did not correlate with the measurements from Session 3. Our results do not reject the possibility that there is in fact a reliable individual differences in the pupil response. Simulations suggest that within-session measurement noise could have obscured a true individual difference effect. Analysis of individual subject data also failed to show a relationship between individual differences in melanopsin function as elicited through the silent substitution and PIPR approaches (Supplementary Figure 4). In future studies, increasing the number of trials and improving pupillometry quality could reduce within-session measurement error and perhaps reveal reproducible individual differences in response.

In Session 3, we examined the effect of a multiplicative increase in stimulus intensity. This manipulation increased the radiance of both the stimulus and the background. For the silent substitution stimuli that targeted either the cones or melanopsin, this change in stimulus intensity did not alter the pupil response. While retinal irradiance was increased in Session 3, the contrast of the silent substitution modulations remained constant at 400%. Therefore, within this stimulus regime, the pupil response to silent substitution stimuli appears to be best characterized in terms of the photoreceptor contrast of the modulation.

These results also allow us to discount the possibility of inadvertent rod stimulation by the melanopsin-directed stimulus. The spectral sensitivity functions of melanopsin and rhodopsin overlap. Consequently, the melanopsin-directed silent substitution stimulus has substantial calculated contrast (~320%) upon the rod photoreceptors. Because the stimulus background is in the photopic range, we generally assume that the rods are saturated, and thus this rod contrast does not contribute to the observed pupil response. This assumption, however, may be challenged by recent work that finds rods can signal above their nominal saturation threshold^48^. It is therefore reassuring to observe in the current study that the pupil response is unchanged with the increased stimulus intensity used in Session 3. If there were a substantial rod contribution to the pupil response measured to the melanopsin-directed stimulus in Sessions 1 and 2, we would expect that this contribution would become smaller at the higher background level. The equivalence of the pupil response suggests that any rod signals are minimal under these conditions.

Conversely, the post illumination pupil response (PIPR) evoked by the chromatic stimuli was enhanced by the increase in stimulus intensity in Session 3. Similar to the silent-substitution stimuli, the photoreceptor contrast produced by the red and blue stimuli is in principle unchanged in Session 3. However, small imperfections in the control of the dim background light levels used for the PIPR stimuli could produce substantial changes in stimulus contrast. While our stimulus measurements indicate fairly consistent calculated contrast between the experimental sessions (Supplementary Table 1), actual variation in the contrasts produced by the PIPR stimuli remains a possible explanation for the enhanced responses to the PIPR stimuli seen in Session 3. It is also possible, however, that the increased pupil response to the PIPR stimuli is a real effect of the change in stimulus intensity. When stimuli are presented against dark backgrounds, changes in intensity can lead to substantial changes in rod activation, which could then alter the response.

We note that our implementation of the PIPR paradigm differs from that used in many other studies, due to the particular nature of our apparatus. For example, the change in stimulus intensity examined in Session 3 increased both the stimulus and background light levels. This is unlike previous studies of the dependence of the PIPR upon intensity^29,49^, in which the background presumably was held fixed across changes in the intensity of the chromatic pulses. Our apparatus also imposes gamut limits that restrict how dark we can make the background and how intense we can make the chromatic pulses. Additionally, many PIPR studies make use of a Ganzfeld dome and thus provide a greater spatial extent of stimulation than used here. These difference likely account for the smaller magnitude of post-illumination pupil response that we obtain in comparison to other studies^18,29,50^. That the PIPR depends on the specifics of the stimuli is an important consideration when comparing results obtained with this paradigm.

Overall, we find that the melanopsin-mediated pupil response at the group level is stable over time, consistent across stimulus conditions, and reflective of known melanopsin physiology. Various clinical conditions, including light sensitivity, may result from an alteration of melanopsin function. Our results suggest that silent-substitution pupillometry can be used to test such hypotheses.

## Supplementary Materials

**Supplementary Figure 1.**
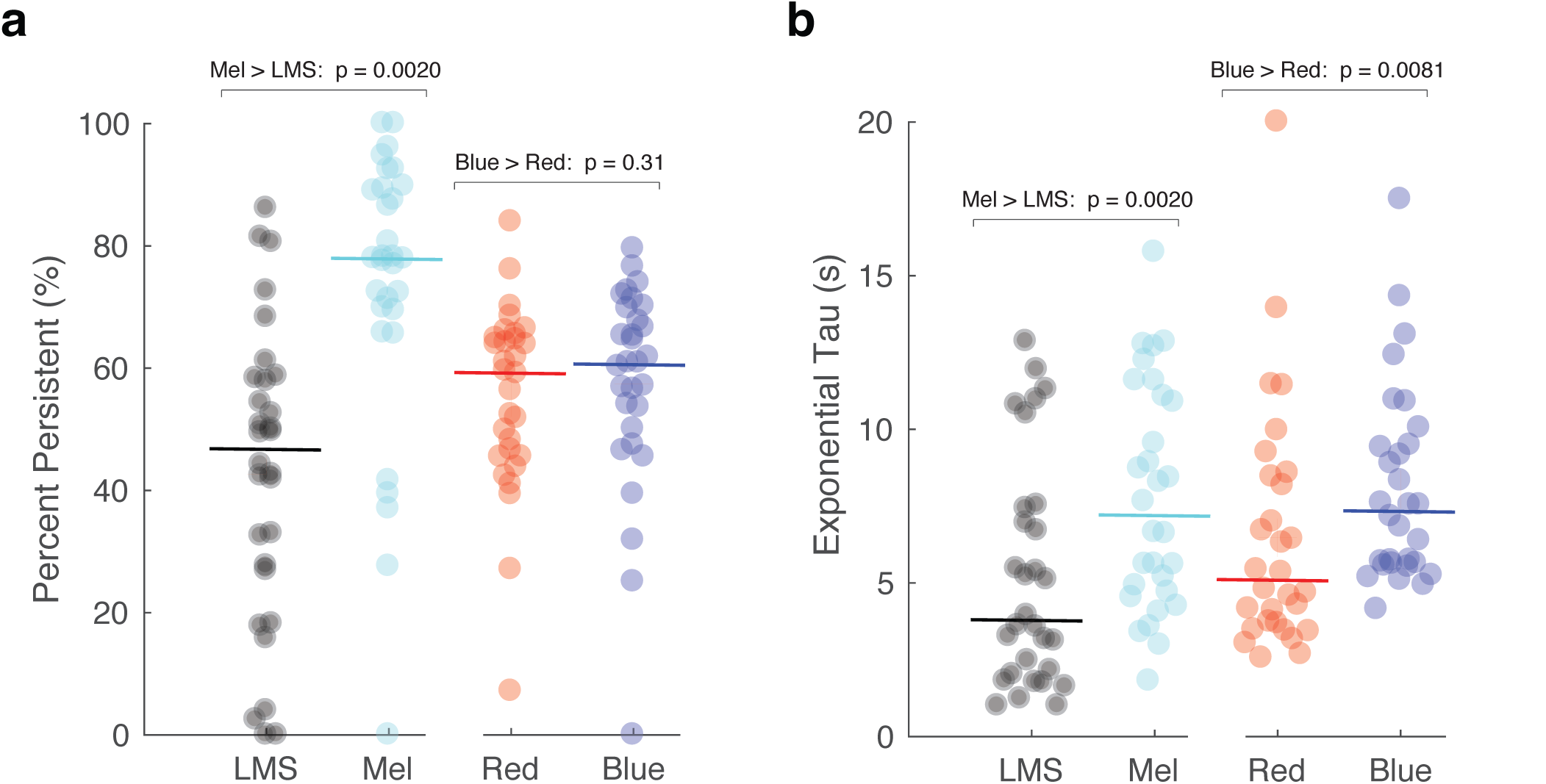
The effect of parameter locking upon model results. Figure 4 presents two model-derived measures (‘percent persistent’ and τ_exponential_) compared between stimuli. This model fitting included other parameters, and it is possible that the observed results reflect a change in these other parameters as opposed to the parameters of interest. To examine this possibility, we re-ran the analysis holding the remaining parameters constant between our compared stimulus conditions. (a) The percent persistent measurement was made for the silent substitution data while fixing the three temporal parameters (τ_exponential_, τ_gamma_, τ_delay_) at the average value across the Mel and LMS response. Similarly, the amplitude of response evoked from the PIPR stimuli was modeled while fixing the temporal parameters at the mean value across the red and blue stimuli. The results of this analysis are largely consistent with the results presented in Figure 4. (b) The complementary analysis, now conducted by holding all model parameters fixed except for τ_exponential_.

**Supplementary Figure 2.**
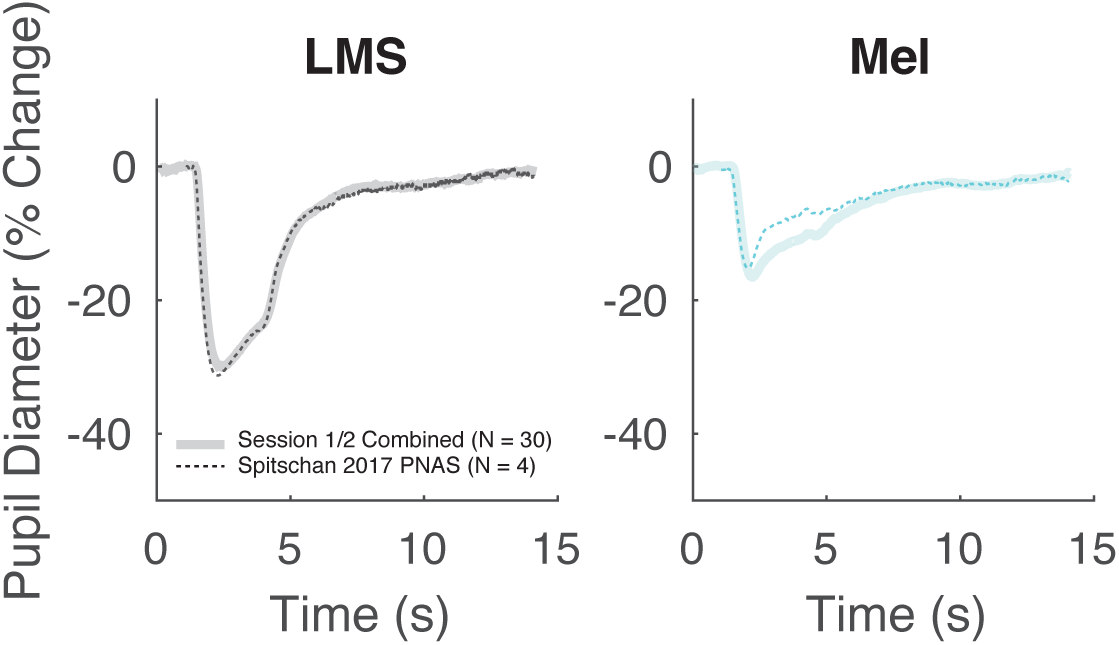
The pupil response evoked by the silent substitution stimuli in Session 3 of the current study was compared to that observed in a prior study of a small number of subjects (N=4). The amplitude and form of response is similar. As compared to the current study, the prior study featured a larger stimulus field (64° vs. 27.5°), but was otherwise similar in stimulus contrast and stimulus background radiance.

**Supplementary Figure 3.**
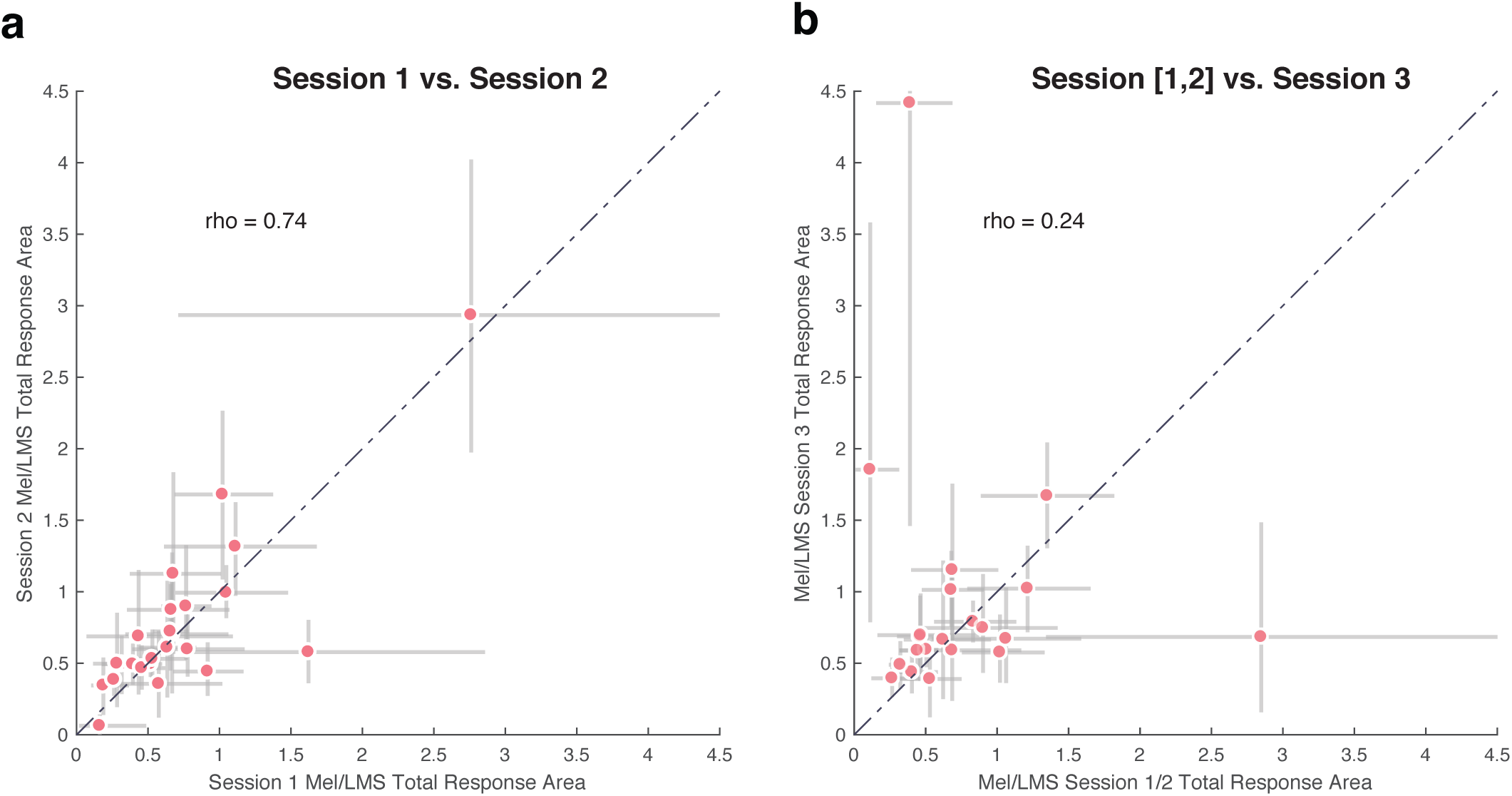
We tested for individual differences in the relative pupil response to the melanopsin‐ and cone-directed stimuli. The total modeled area of pupil response to the melanopsin and cone stimulus was obtained and expressed as a ratio. (a) When compared between Sessions 1 and 2, individual differences in response ratio were well reproduced (Spearman’s rho = 0.74, N = 24 subjects). (b) The same analysis, now comparing the average measurement from Sessions 1 and 2 with the measurement from Session 3 (Spearman’s rho = 0.24, N = 21 subjects). A weaker correlation was seen. The error bars reflect the 10-90% confidence interval, obtained via bootstrap analysis across trials within a subject. As the magnitude of these error bars are large relative to the variation across subjects, it is possible that within-session measurement noise limits our ability to detect if there is in fact a stable individual difference in relative pupil response to melanopsin and cone stimulation.

**Supplementary Figure 4.**
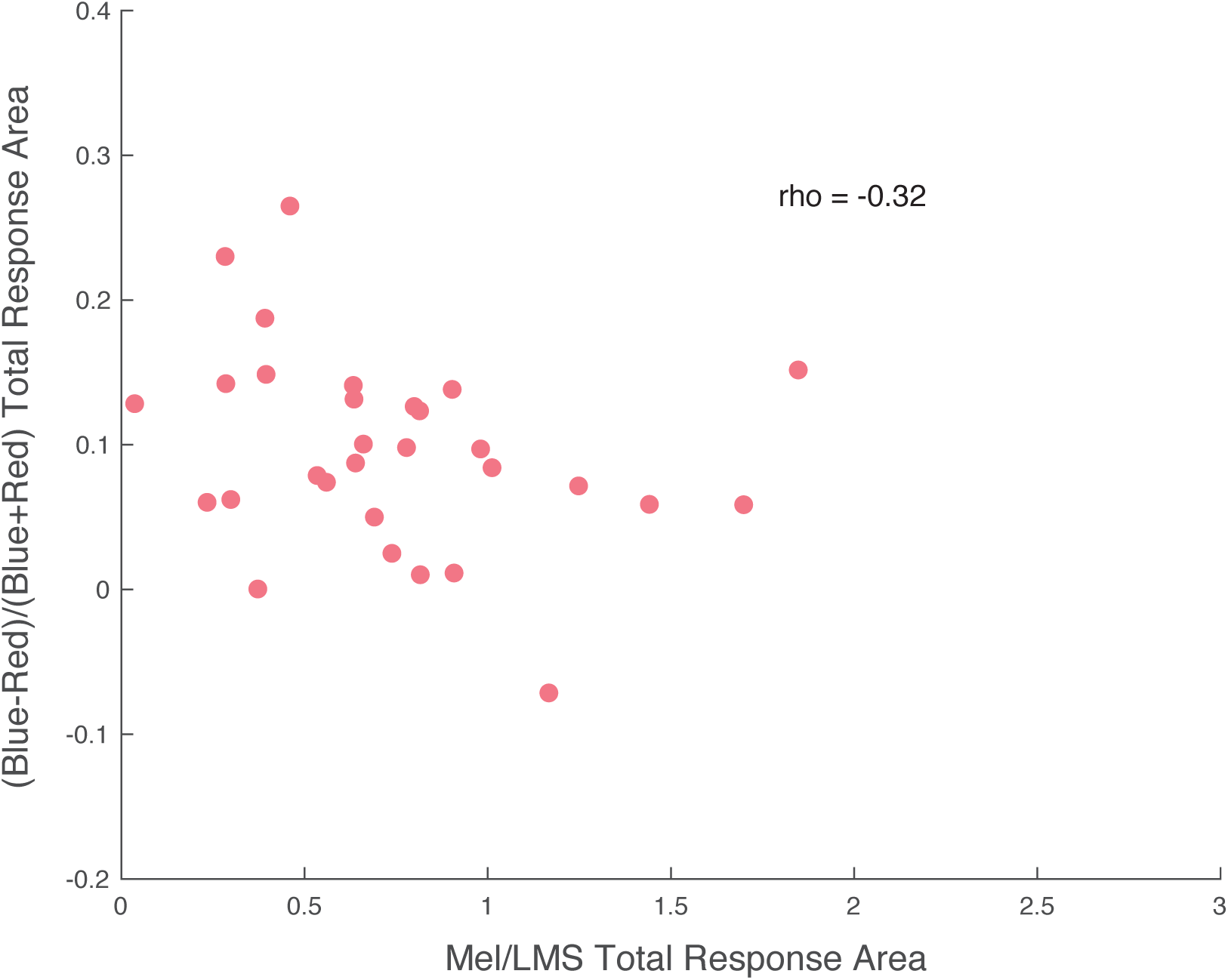
A test for individual differences in melanopsin function as elicited by the silent substitution and PIPR approaches. We fit the three-component model to the average response for each subject to all trials of each stimulus type across Sessions 1 and 2. The PIPR effect was expressed as the difference in the area of response to the blue and red stimuli, divided by the sum of the response areas. This normalization was needed to account for individual differences in overall pupil responses, independent of variation in melanopsin sensitivity per se. The melanopsin effect in the response to the silent substitution stimuli was expressed as the ratio of the response area for the melanopsin-directed stimulus divided by that evoked by the cone-directed stimulus. While a correlation with a positive slope would be expected if these two measurements reflect an underlying individual difference in melanopsin sensitivity, this was not observed (Spearman’s rho = ‐0.28).

**Supplementary Table 1.**
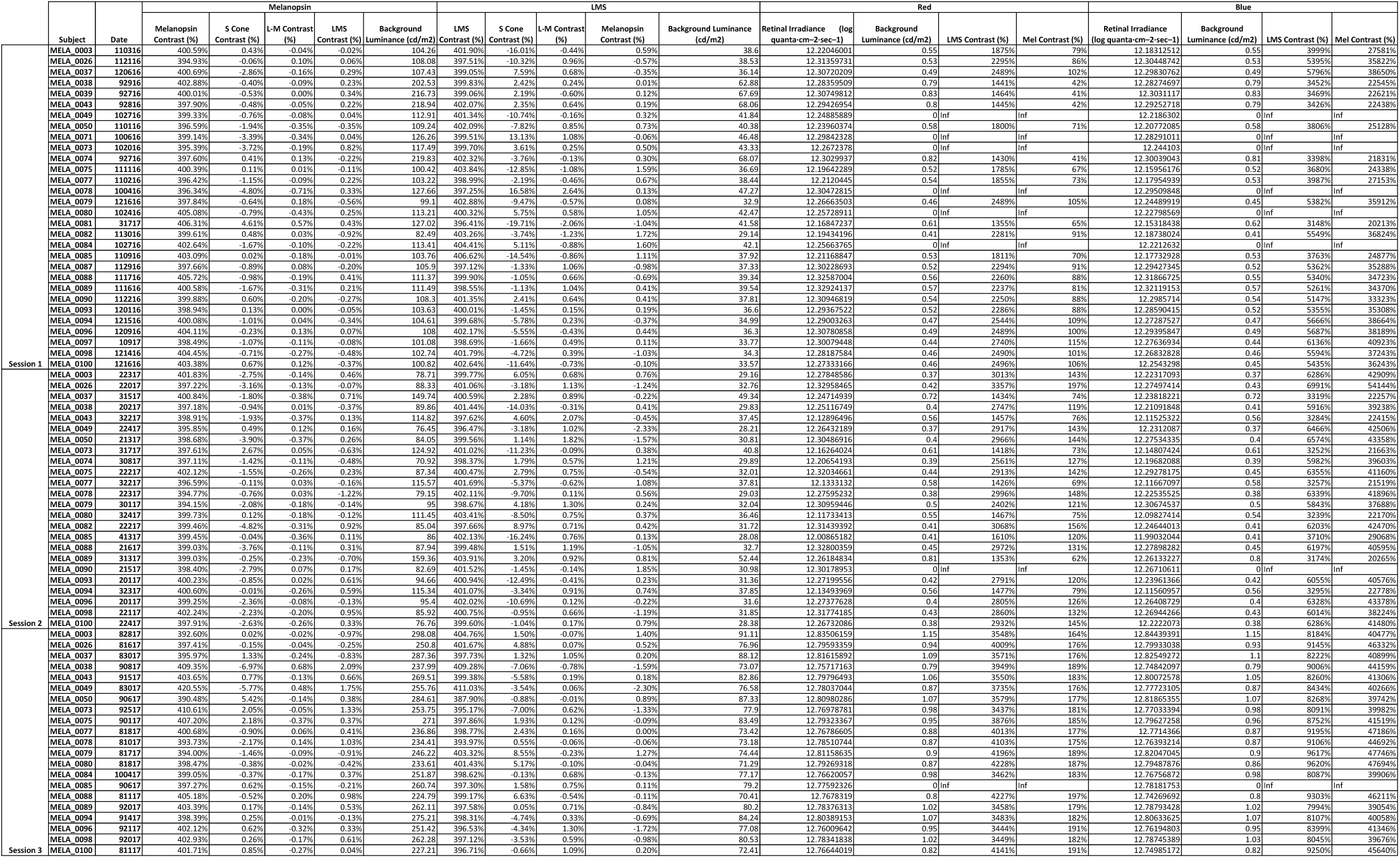
Summary of validation measurements for each subject for each session. For each subject, for each session, 5 measurements of the spectral power distribution of each stimulus type are taken before and after the experiment. This table summarizes the median background luminance and contrast both on the targeted photoreceptor mechanism, as well as the inadvertent splatter on receptors nominally isolated) for each silent substitution stimulus, as well as the median retinal irradiance and background luminance for each PIPR stimuli. For several subjects, the median calculated photoreceptor contrast for the PIPR stimuli was infinite (‘Inf’); this value was obtained when the background light level was below the intensity floor that could be measured with our radiometer.

**Supplementary Table 2.** Median model parameters for each stimulus by session. We fit a three-component model to the average pupil response for each subject for each session to each stimulus. This table shows the median value across all subjects for each parameter of the model.

| **LMS** | *Transient Amplitude*<br>*(% change \* s)* | *Sustained Amplitude*<br>*(% change \* s)* | *Persistent Amplitude*<br>*(% change \* s)* | *Exponential Tau*<br>*(s)* | *Gamma Tau*<br>*(ms)* | *Delay*<br>*(ms)* |
| --- | --- | --- | --- | --- | --- | --- |
| Session 1 | -15.10 | -34.18 | -59.96 | 3.77 | 393.91 | -256.37 |
| Session 2 | -14.99 | -30.06 | -57.24 | 2.74 | 393.48 | -269.45 |
| Session 3 | -16.87 | -37.42 | -61.01 | 3.37 | 380.52 | -265.58 |
| **Mel** | *Transient Amplitude*<br>*(% change \* s)* | *Sustained Amplitude*<br>*(% change \* s)* | *Persistent Amplitude*<br>*(% change \* s)* | *Exponential Tau*<br>*(s)* | *Gamma Tau*<br>*(ms)* | *Delay*<br>*(ms)* |
| Session 1 | -9.30 | 0.00 | -61.34 | 5.49 | 349.21 | -375.84 |
| Session 2 | -9.67 | -0.53 | -48.78 | 5.52 | 329.82 | -399.72 |
| Session 3 | -9.07 | -1.74 | -54.97 | 5.96 | 335.41 | -367.05 |

| **Red** | *Transient Amplitude*<br>*(% change \* s)* | *Sustained Amplitude*<br>*(% change \* s)* | *Persistent Amplitude*<br>*(% change \* s)* | *Exponential Tau*<br>*(s)* | *Gamma Tau*<br>*(ms)* | *Delay*<br>*(ms)* |
| --- | --- | --- | --- | --- | --- | --- |
| Session 1 | -21.91 | -41.98 | -112.87 | 5.08 | 468.93 | -213.95 |
| Session 2 | -24.70 | -39.04 | -118.25 | 4.55 | 501.50 | -209.22 |
| Session 3 | -27.31 | -35.80 | -125.38 | 4.01 | 516.65 | -181.37 |
| **Blue** | *Transient Amplitude*<br>*(% change \* s)* | *Sustained Amplitude*<br>*(% change \* s)* | *Persistent Amplitude*<br>*(% change \* s)* | *Exponential Tau*<br>*(s)* | *Gamma Tau*<br>*(ms)* | *Delay*<br>*(ms)* |
| Session 1 | -28.62 | -62.05 | -118.70 | 6.80 | 561.98 | -162.61 |
| Session 2 | -28.87 | -55.11 | -120.35 | 6.57 | 515.79 | -184.96 |
| Session 3 | -31.54 | -62.27 | -139.37 | 7.89 | 562.46 | -143.06 |

